# hichipper: A preprocessing pipeline for assessing library quality and DNA loops from HiChIP data

**DOI:** 10.1101/192302

**Authors:** Caleb A. Lareau, Martin J. Aryee

## Abstract

Mumbach *et al.* recently described HiChIP, a novel protein-mediated chromatin conformation assay that lowers cellular input requirements while simultaneously increasing the yield of informative reads compared to previous methods (1). To facilitate the dissemination and adoption of this assay, we introduce **hichipper** (http://aryeelab.org/hichipper), an open-source HiChIP data preprocessing tool, with features that include bias-corrected peak calling, library quality control, DNA loop calling, and output of processed data for downstream analysis and visualization (Figure 1a).

**Figure 1:**
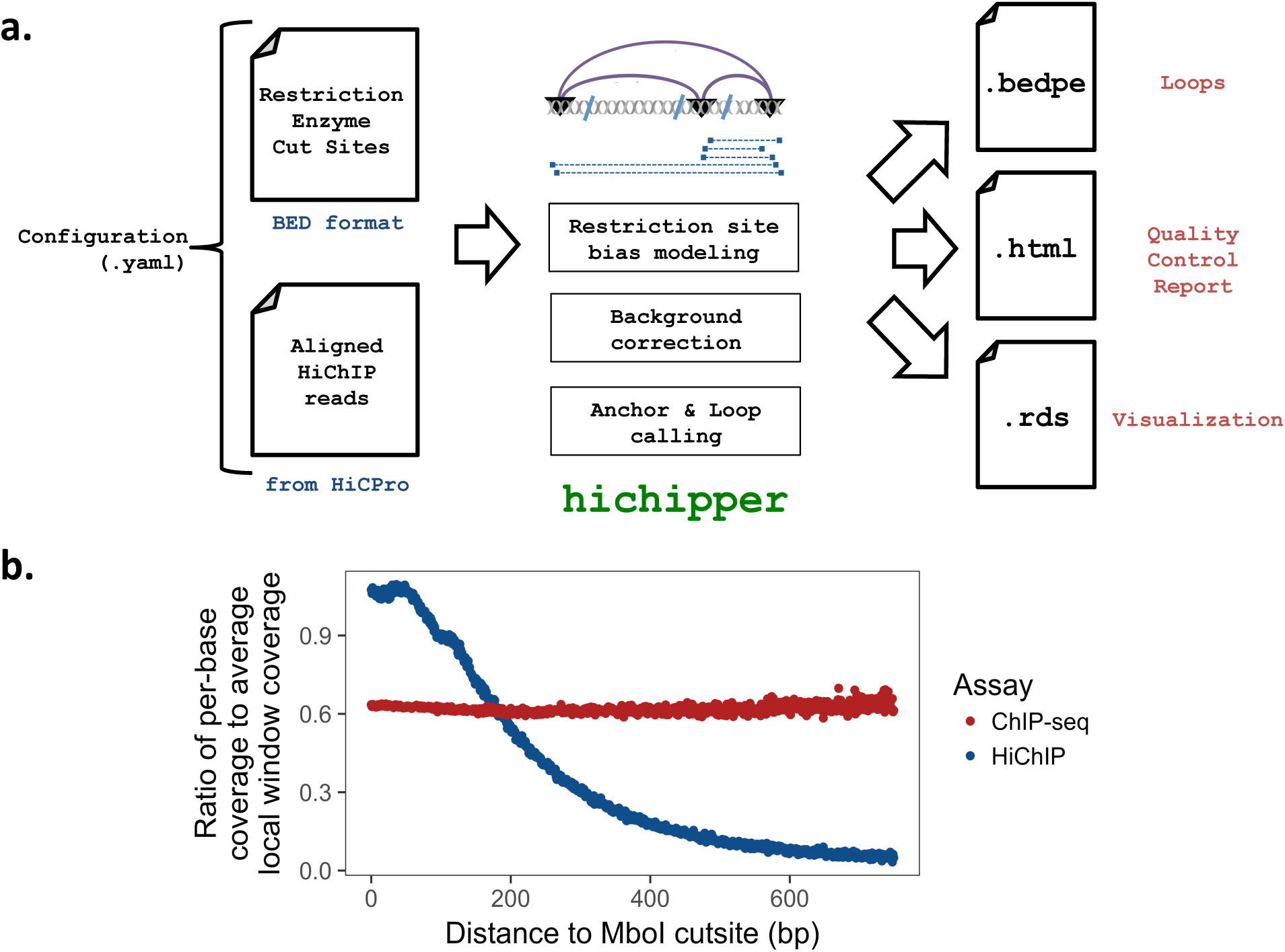
a) Overview of the **hichipper** analysis pipeline. **hichipper** requires aligned and annotated in-teraction files from preprocessing tools such as Hi-C Pro (2) as well as a .bed file of restriction sites. The **hichipper** pipeline identifies loop anchors using a background model that accounts for the effect of restriction site proximity on read density. The output of **hichipper** can be used for quality control, visualization, and downstream topology analysis of HiChIP data. b) Ratio of per-base coverage to local MACS-estimated local window background signal as a function of distance to nearest MboI cutsite for a HiChIP sample (blue) and a ChIP-seq sample (red). Both samples represent published mouse ESC with SMC1 (cohesin) ChIP.

---

The HiChIP protocol, unlike assays such as ChIA-PET and ChIP-seq, involves a restriction enzyme treatment step. This results in a read density landscape that is biased by proximity to restriction sites (Figure 1b), hampering identification of peaks representing DNA loop anchors and hence loops. **hichipper** implements a HiChIP-specific background model that accounts for this read density bias by modeling the background read density as a function of proximity to restriction sites, resulting in significant improvements in identification of loop anchors and overall loop calling. Further, we have found that taking the location of restriction enzyme cut sites into account also boosts sensitivity by increasing the fraction of reads that can be assigned to loops. **hichipper** outperforms existing non-HiChIP specific tools in terms of identifying long-range (> 100kb) loops with a typical anchors resolved at a 2.5kb resolution (**Supplemental Methods**).

Additionally, **hichipper** produces a quality control report that allows evaluation of library preparation, including factors such as the efficiency of proximity ligation and chromatin immunoprecipitation. We provide example quality control reports from both successful and unsuccessful HiChIP experiments to guide new users in assessing their own library quality.

The **hichipper** package, documentation and QC report examples are available online at http://aryeelab.org/hichipper. Details of the background model and implementation are available in the **Supplemental Methods**.

## Supplementary Information for **hichipper**

### Highlights

- HiChIP read distribution is biased by proximity to restriction enzyme cut sites, hindering the identification of true loop anchors.
- `hichipper` employs a background model that incorporates the effect of restriction site bias when identifying loops.
- Loop anchors called using the `hichipper` background model are enriched for overlap with ChIP-seq peaks and promoters compared to those identified by standard approaches.
- Loops identified by `hichipper` are more sensitive to putative long range interactions and can define loops at higher resolution than other methods.

### Data

To characterize the unique properties of HiChIP, we compared published data from SMC1 ChIP-seq,^1^ ChIA-PET,^2^ and HiChIP^3^ experiments. Specifically, the samples that were used for the primary comparison are available from the Sequence Read Archive (SRA) under the following accession numbers: HiChIP: SRR3467179; ChIA-PET: SRR1296617; ChIP-seq: SRR058981, SRR058982. For consistency, all samples were aligned to the mm9 reference genome using Bowtie2^4^ after modifying the reads as appropriate to the specific assay (*i.e.* linker cutting in ChIA-PET through Mango^5^; restriction enzyme ligation cutting in HiChIP through HiC-Pro^6^). Additional data described in the original HiChIP paper^3^ were processed similarly. Peaks were called using MACS2^8^ (`-q 0.01 --nomodel --extsize 147`).

### Restriction site bias influences HiChIP read distribution

After examining HiChIP read distributions we noted key differences relative to those from ChIA-PET and ChIP-seq. Supplemental Figure 1 depicts read pileups for two different genomic loci in mouse embryonic stem cells (mESC). Since HiChIP involves a restriction enzyme treatment step (like HiC), we also display occurrences of the MboI retriction site motif (`GATC`) in the bottom track indicated by blue vertical lines. While ChIA-PET reads have a distribution that resembles that of the ChIP-seq track, HiChIP shows a background read distribution that appears to be biased by proximity to MboI motifs. Likely as a result of this, MACS identifies vastly more peaks from the HiChIP sample (186,071) compared to the number expected from ChIP-seq (65,204).

As an alternative approach to HiChIP peak calling, Mumbach *et al.* suggest using only dangling-end and self-ligation reads (denoted as “self” for the rest of this document). We find that this approach is effective in reducing background signal (Supplemental Figure 2), but it also appears to reduce the sensitivity of anchor identification as the number of peaks called by MACS (24,870) is less than half that expected from ChIP-seq in these datasets. The number of peaks called using different classes of input reads is summarized in Supplemental Figure 3. In particular, the plot shows the number of anchors identified for 1) ChIA-PET, 2) ChIP-seq, 3) HiChIP using all reads, and 4) HiChIP using only self-ligation and dangling-end reads as suggested in Mumbach *et al.*

**Supplemental Figure 1:**
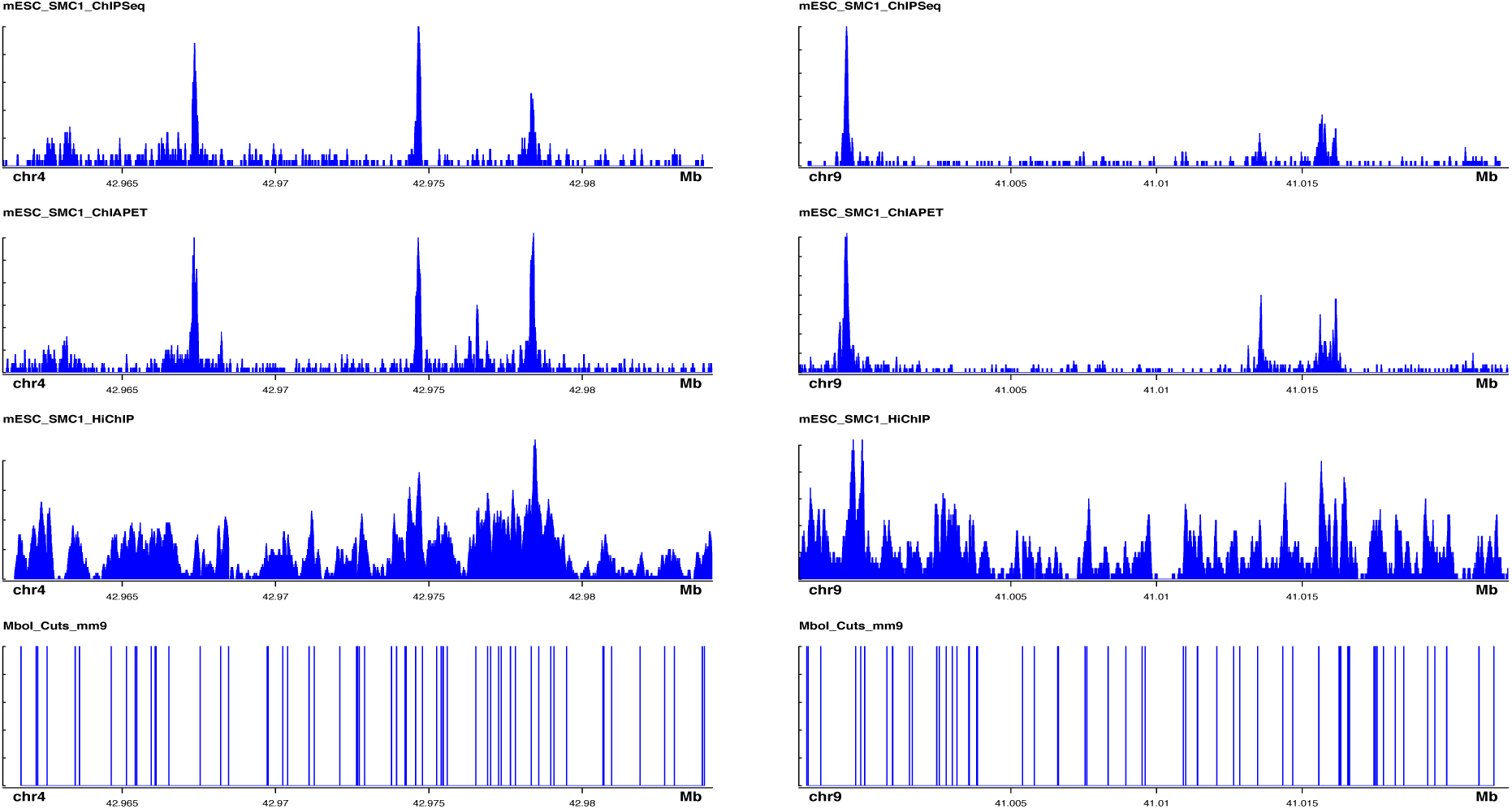
Total read pileup distributions targeting mESC SMC1 across two different genomic regions. Occurrences of the MboI motif in mm9 are shown in the bottom track and indicated by a blue vertical line. While ChIP-seq and ChIA-PET generally have similar peak landscapes, HiChIP read density is biased by proximity to restriction sites. This bias leads, in this case, to an inflated number of identified peaks when using standard peak callers.

**Supplemental Figure 2:**
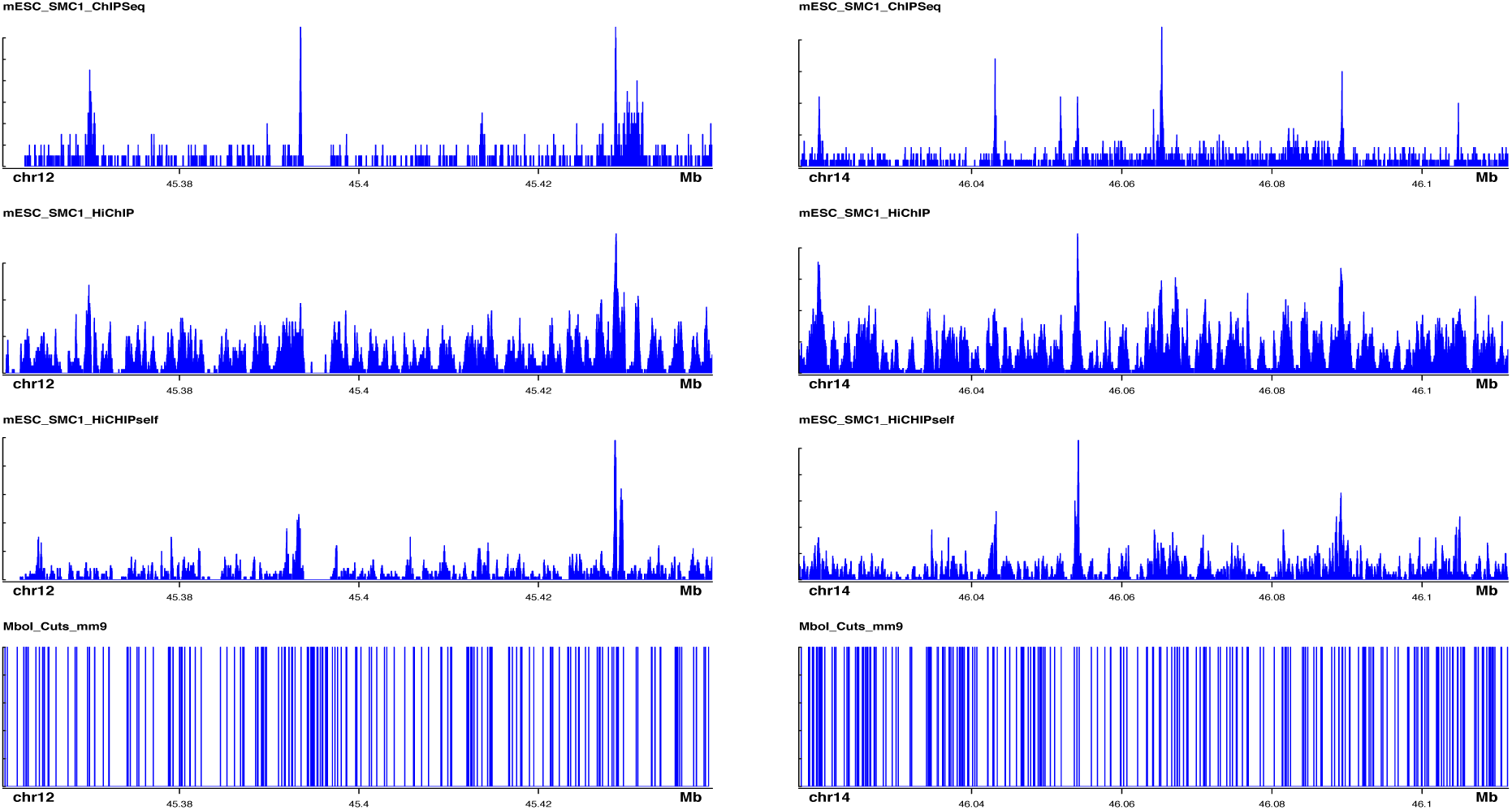
Read pileup distributions for two additional genomic loci showing a new track for only the self-ligation reads. While using only the self-ligation reads removes a considerable proportion of the background and limits the number of peaks called, the reduction in read count reduces power to detect loop anchors.

**Supplemental Figure 3:**
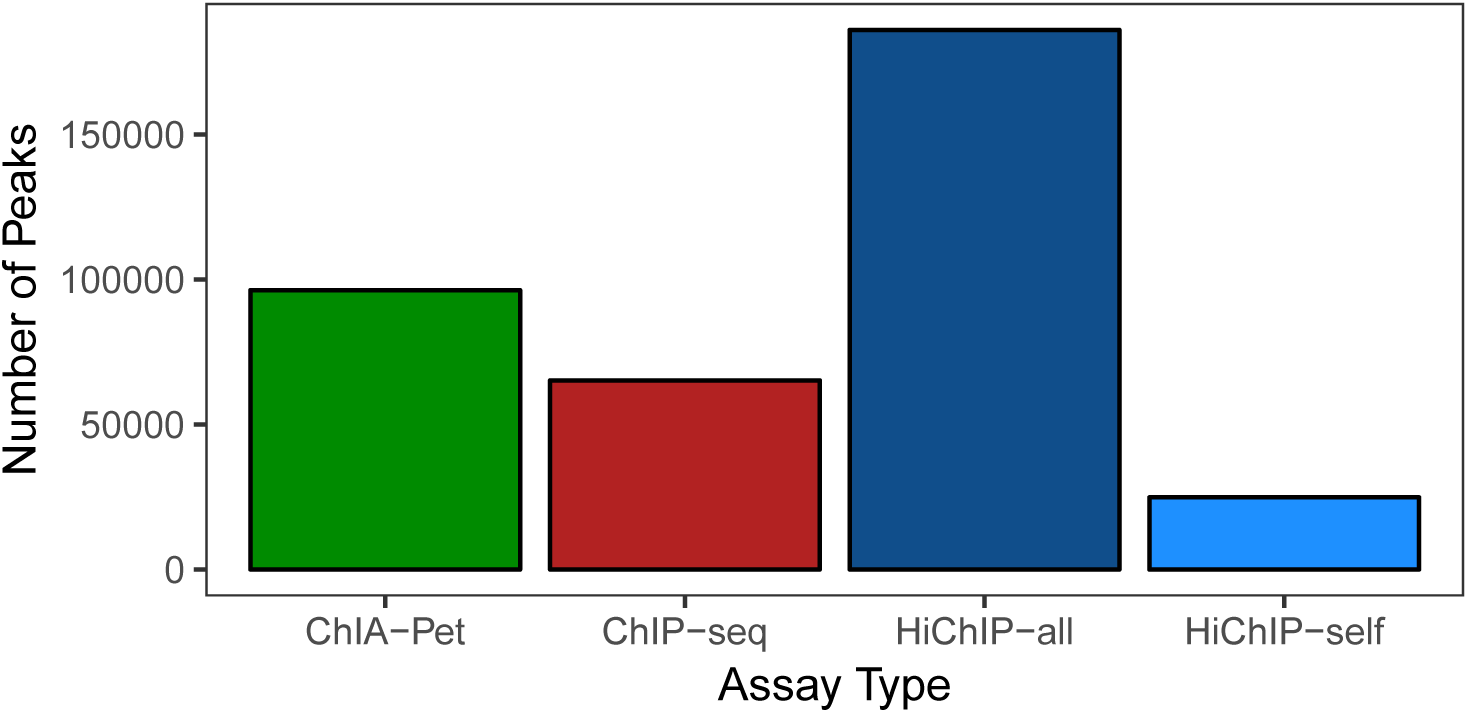
Number of peaks called at FDR = 0.01 for ChIA-PET, ChIP-seq and HiChIP using MACS2. For HiChIP, peaks were called using all reads as is conventional in most preprocessing pipelines (dark blue) and separately using only self-ligation and dangling-end reads as suggested in Mumbach *et al.* (light blue)

In brief, calling HiChIP peaks using all reads (Supplemental Figure 3, dark blue) as would be standard in ChIA-PET preprocessing pipelines leads to a several-fold increase in peak calls relative to ChIP-seq (red). Conversely, using only self-ligation and dangling-end reads (light blue) can result in low sensitivity by calling too few peaks. Moreover, by definition, the filtering criterion of using only self-ligation reads removes all paired-end reads that could support interactions, which may lead to genomic loci associated with looping to be missed in anchor inference. Thus, we sought to define an algorithm that 1) uses all reads from HiChIP and 2) explicitly models the bias associated with proximity to restriction enzyme cut sites to call loop anchor peaks.

### hichipper peak calling

When MACS2^8^ determines peaks from sequencing data, the algorithm identifies regions where the read pileup is sufficiently higher than a conservative estimate of the background read density, estimated either from an independent control sample or from the ChIP sample of interest itself. Specifically, MACS2 estimates *λ*_BG_, the genome-wide average read density. Additionally, the software computes per-peak small (*λ*_1*K*_) and large (*λ*_10*K*_) background parameters representing the read density within 1kb and 10kb around each putative peak. The parameter used to model the background read density, *λ*_local_, is conservatively computed as

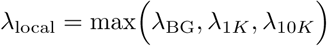

Per-peak p-values and q-values are computed under the null hypothesis that the observed read coverage count at a putative peak is generated from a Poisson distribution parameterized by *λ*_local_. While this background read density estimation method works well for ChIP-seq and ChIA-PET data, the clear bias of read density related to restriction cut site proximity in HiChIP suggests that a different choice of background model for peak (*i.e.* loop anchor) calling may be more appropriate for this assay.

To characterize the restriction site bias of HiChIP signal, we examined the ratio of read coverage signal to *λ*_local_ as a function of distance to the nearest MboI cutsite (Supplemental Figure 4). In ChIP-seq (red) where no restriction enzyme is used, the ratio is unaffected by cut site proximity as would be expected. In contrast, a characteristic trend emerges in HiChIP data where regions of the genome close to a restriction enzyme cut site have considerably more signal, and thus a higher likelihood of being identified as a peak, than regions distant from restriction sites. We found that this relationship was present in all mESC SMC1 HiChIP replicates (Supplemental Figure 5) and was even more extreme in the GM12878 SMC1 replicates (Supplemental Figure 6).

**Supplemental Figure 4:**
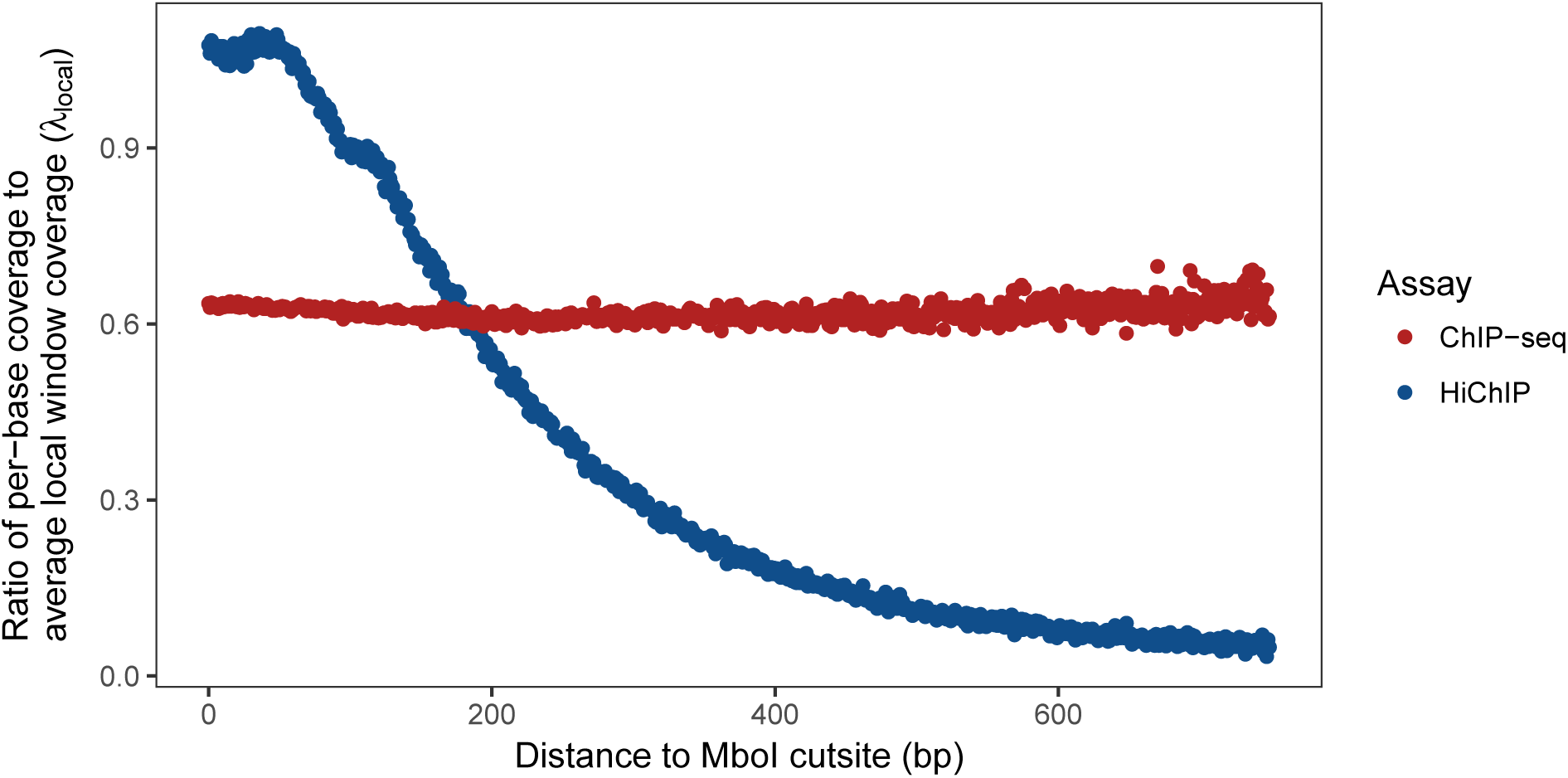
Ratio of per-base coverage to local MACS-estimated local window background signal as a function of distance to nearest MboI cutsite for a HiChIP sample (blue) and a ChIP-seq sample (red). The trend observed in the HiChIP sample reveals a mis-specified background. As a consequence, an inflated number of peaks near MboI cut sites are called from the HiChIP data while putative peaks far from cut sites are underrepresented. Both samples represent mouse ESC with SMC1 (cohesin) ChIP.

**Supplemental Figure 5:**
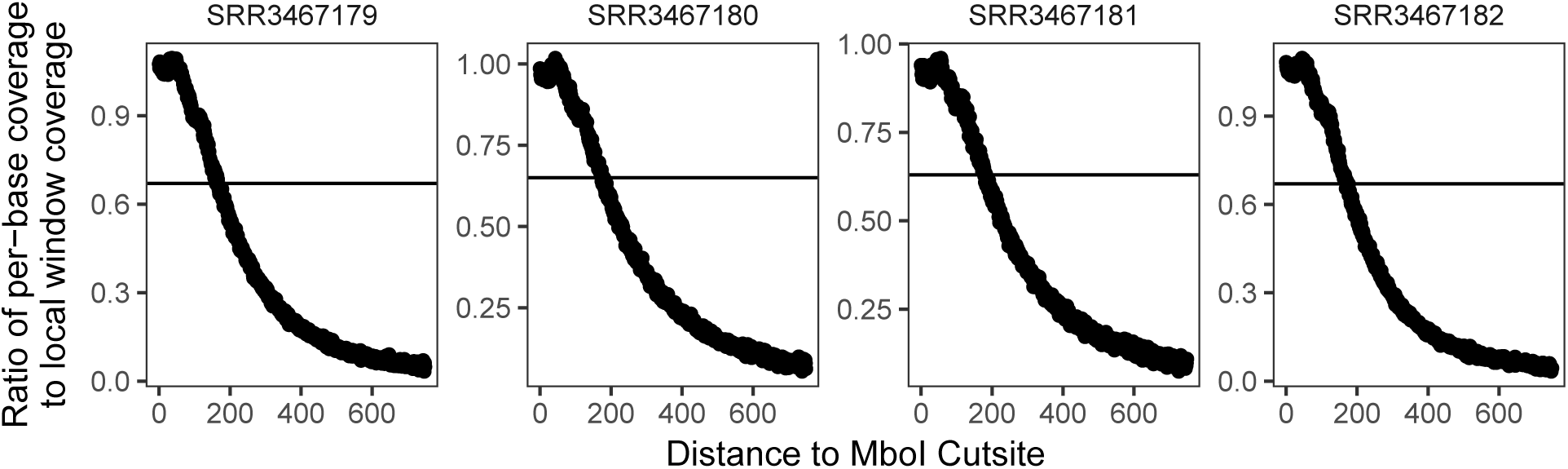
Ratio of per-base coverage to local MACS-estimated background signal as a function of distance to nearest MboI cutsite for four mESC SMC1 HiChIP Samples. The black horizontal line represents the per-sample global mean.

**Supplemental Figure 6:**
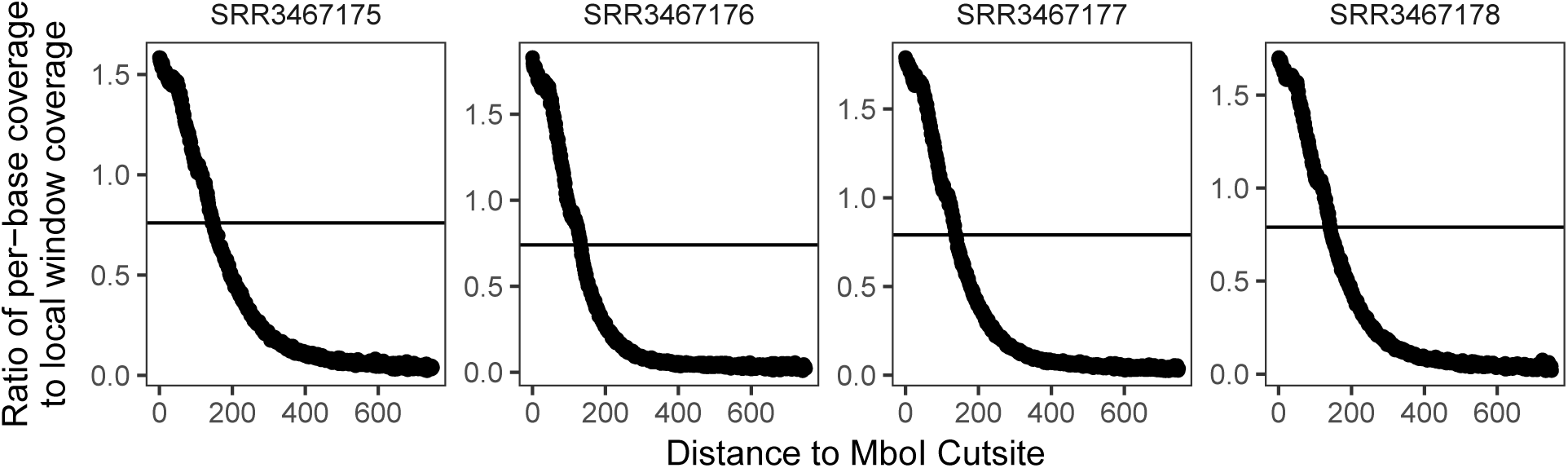
Ratio of per-base coverage to local MACS-estimated background signal as a function of distance to nearest MboI cutsite for all four GM12878 SMC1 HiChIP Samples. The black horizontal line represents the per-sample global mean.

To model the restriction site bias when performing peak calling, we modify the parameterization of the background signal Poisson model implemented in MACS2. In particular, we fit a smoothing spline, *f* (*d*), (per sample) to the curve shown in Supplemental Figure 4–6 and compute a modified background parameter 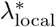 as a function of the distance *d* of the midpoint of a putative peak to its nearest restriction fragment cut site. In line with the conservative nature of the MACS2 peak calling, we define 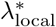 as

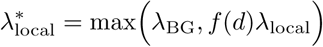

`hichipper` then uses this restriction site distance-dependent 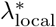as the parameter for the background Poisson model when computing per-peak p-values and q-values. In effect, the modified Poisson model reduces the number of peaks called near restriction enzyme cut sites while simultaneously making regions far from cut sites more likely to be called peaks for a given read density. This specification retains the conservative implementation of the MACS2 peak calling (via setting a floor of *λ*_BG_) while simultaneously increasing the stringency of peak calling near MboI sites and relaxing the stringency at genomic loci far from MboI sites.

### Evaluation of hichipper peak calling

To evaluate the performance of the modified background model, we called peaks with and without the distance-dependent background correction implemented in `hichipper` (referred to as the “hichipper background model”) using published mESC SMC1 HiChIP data. The `hichipper` background model identifies approximately the same number of peaks as we would expect from SMC1 ChIP-seq data (indicated by the dashed black line) whereas the standard MACS background model results in an inflated number of peaks (Supplemental Figure 7).

**Supplemental Figure 7:**
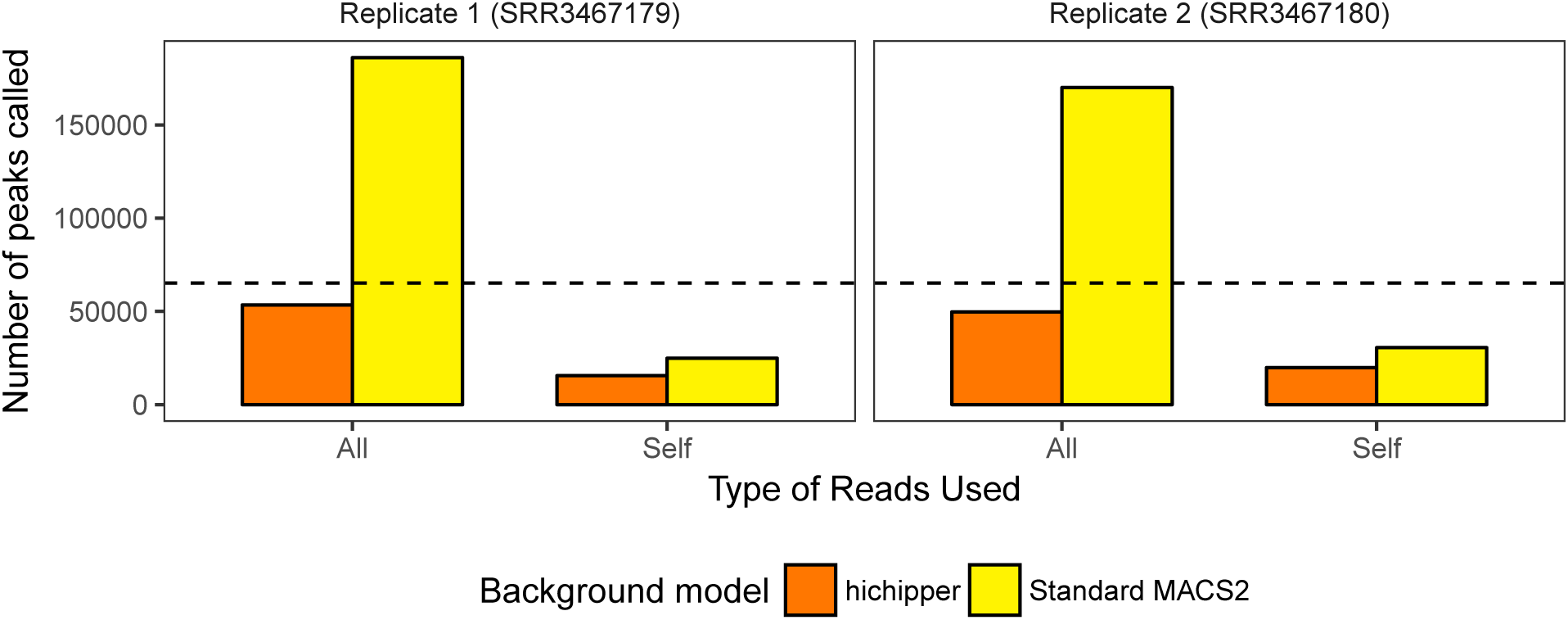
Numbers of peaks called for mESC SMC1 HiChIP replicates using two differerent background models. The dotted black line represents the number of peaks called in an SMC1 ChIP-Seq sample. The x-axis shows the class of reads used to call peaks (all or self-ligation only) with the background model used indicated by color.

Of the 53,446 peaks called by `hichipper` using the modified background correction, 29,220 (55%) could be classified as “ChIP-seq validated” (defined as being within 1kb of an mESC SMC1 ChIP-seq peak). The proportion of peaks that could be validated by ChIP-seq is summarised in (Supplemental Figure 8). Of the > 133, 000 peaks that were eliminated by switching from the standard background model to the hichipper background model, only 26.1% were within 1kb of a ChIP-seq peak, suggesting a higher false positive rate in the standard background model set. Moreover, the number the peaks within 1kb of a RefSeq transcription start site increased from 14% (standard MACS2) to 20% (hichipper) using all reads, further suggesting that the modified background correction enriches for transcriptionally relevant loops.

We also noted that hichipper peaks called using only “self” reads had a very high 94% overlap with ChIP-seq peaks (Supplemental Figure 8), with the important caveat that the number of peaks is very low (15,516; Supplemental Figure 7) resulting in a high false negative rate. It is possible that the poor sensitivity obtained with self-only reads would be improved with considerably greater sequencing depth. Overall, regardless of the read input type, our results suggest that the restriction site-aware background model implemented in hichipper improves the concordance of HiChIP and ChIP-seq peak loci.

**Supplemental Figure 8:**
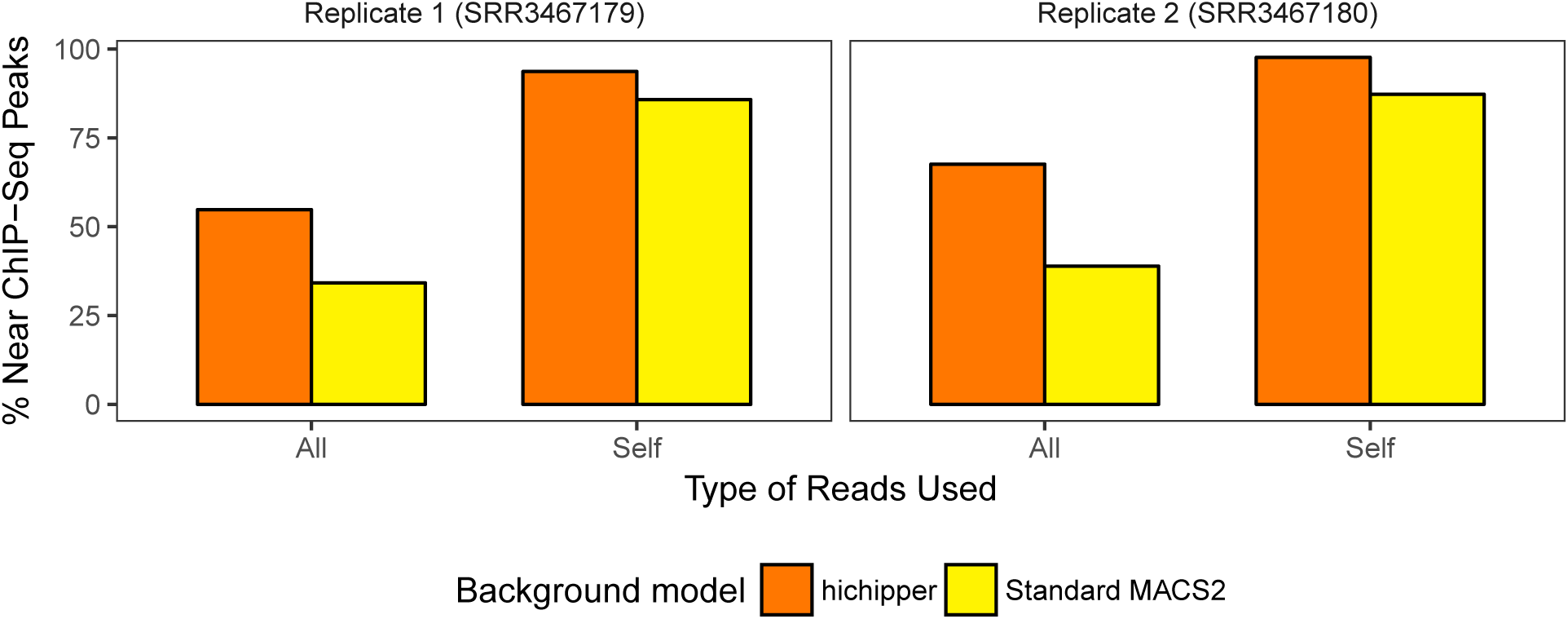
Summary of proportion of peaks within 1kb of an SMC1 ChIP-Seq peak. The restriciton cut site distance-dependent background model implemented in hichipper performs better than the standard model used in MACS2 for enriching with ChIP-seq peaks in each setting examined.

We next sought to characterize `hichipper`-identifed peaks in terms of their involvement in loops. **We define loops as interactions supported by two or more PETs and with a Mango**^5^ **q-value** < 0.01 (see **Loop calling** section), and found that a high proportion (48.7%) of loops involved two ChIP-seq validated loop anchors. In contrast, only 10.8% of the loops were between two non-ChIPseq peaks. These results suggest that HiChIP peaks that are also supported by ChIP-seq are more likely to represent true loop anchors than would be expected from the proportion of ChIP-seq peak/HiChIP anchor overlap (Supplemental Figure 9). As an alternative to peak identification from HiChIP data, hichipper optionally enables users to supply a high-confidence peak/anchor set derived from ChIP-seq or another source. The *a priori*-defined peaks can be specified in the .yaml configuration file.

**Supplemental Figure 9:**
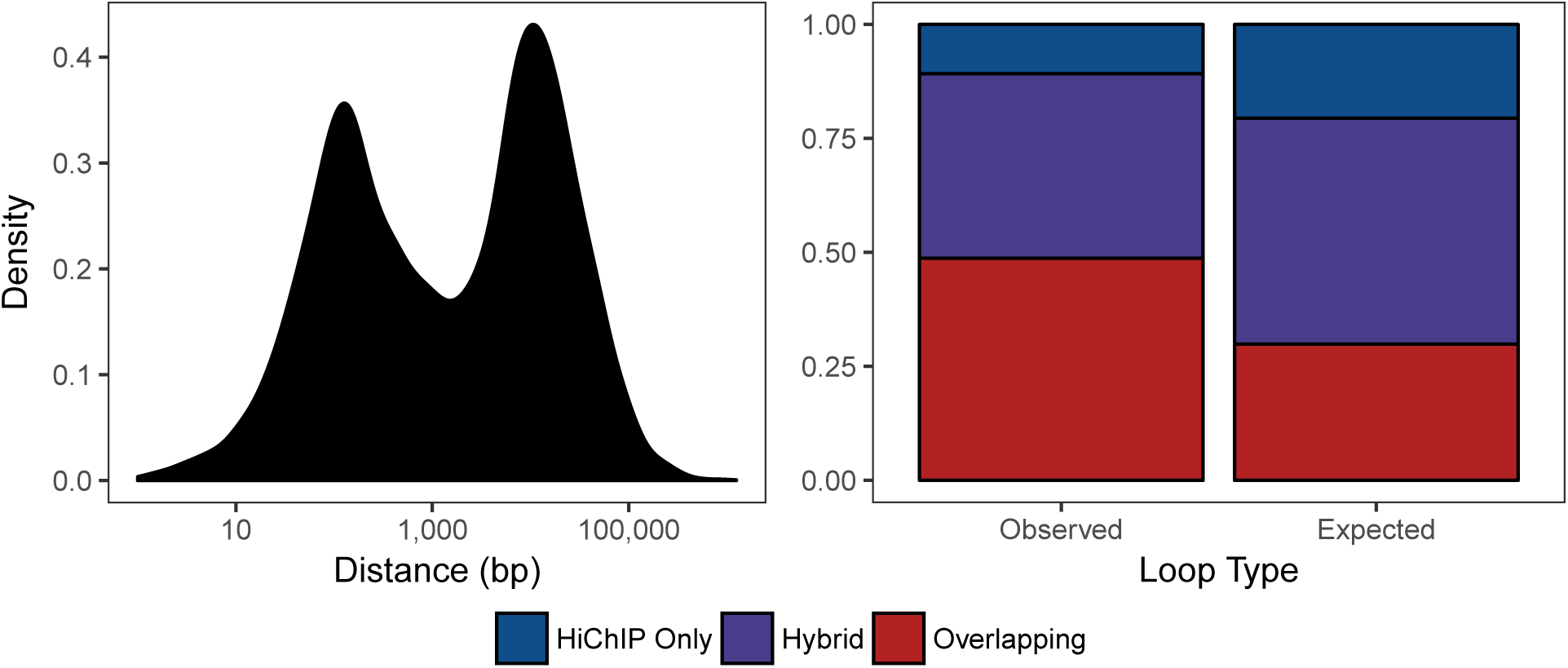
(Left) Distribution of distances between HiChIP-defined anchors and the nearest ChIP-seq peak. (Right) Proportion of putative interactions and PETs called from HiChIP data annotated by overlap with ChIP-Seq peaks. We observe that nearly 50 percent of loops and PETs involve anchors where both overlap ChIP-seq peaks (red).

### Restriction-fragment aware anchor definitions

The use of restriction enzyme fragmentation in HiChIP limits the highest theoretical resolution, as in Hi-C, to the restriction fragment length. Further analysis of peaks identified from ChIP-seq but missed in HiChIP revealed instances where a ChIP-seq peak is positioned near the middle of a long restriction fragment with no nearby HiChIP read density due to the large distance to the closest restriction site (Supplemental Figure 10). HiChIP PETs supporting interactions between anchors such as these tend to localize primarily at the edges of the restriction fragments. To account for this effect, `hichipper` defines loop anchor loci by expanding peaks to the ends of the overlapping restriction fragment(s) as depicted in Supplemental Figure 11. By default, `hichipper` first pads peaks by a fixed window (*i.e.* 500bp) to account for uncertainty in the peak calling (in line with similar preprocessing algorithms) before extending to the restriction fragment edges.

**Supplemental Figure 10:**
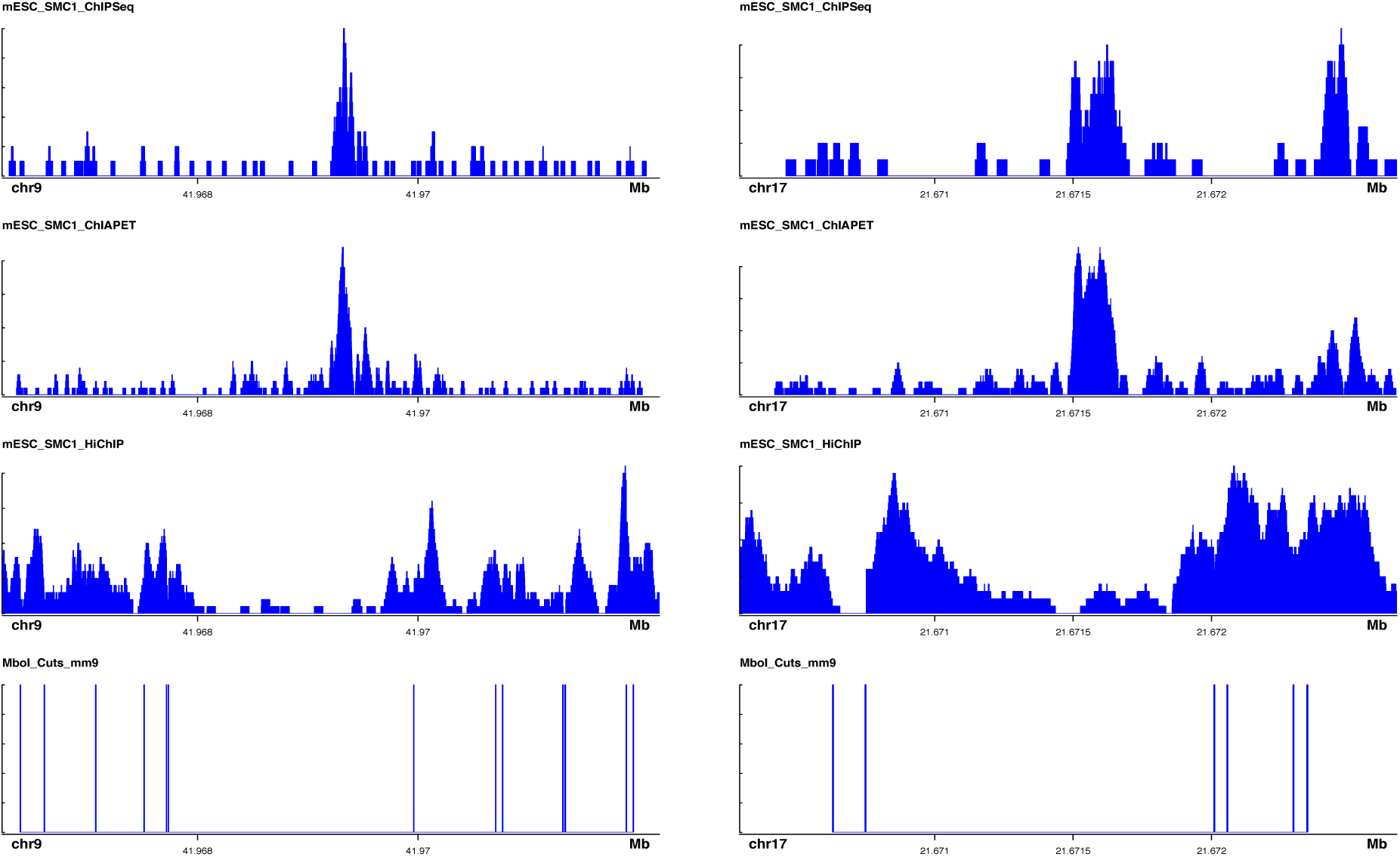
Read pileup distributions for two genomic loci showing a missing peak in HiChIP likely due to the distance spanning the peak and the nearest restriction enzyme motifs. Read density localizes near the edges of the restriction fragment containing the SMC1 peak.

**Supplemental Figure 11:**
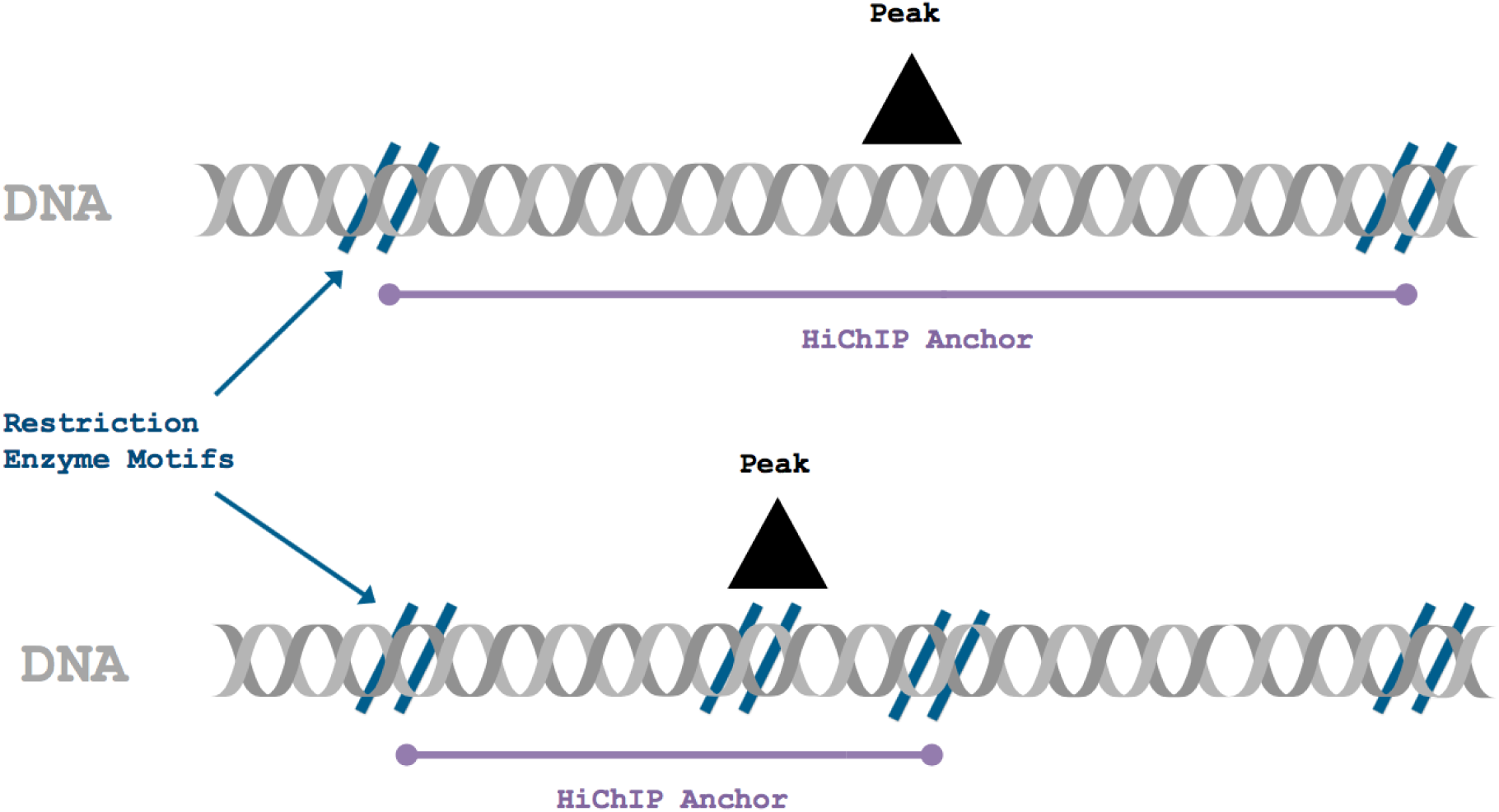
Overview of restriction fragment-aware anchor calling. For a peak that sits on a single restriction fragment (top), `hichipper` extends the peak to the edges of the restriction fragment. When the peak spans one or more restriction fragments (bottom), the anchor is extended to include the entire length of overlapping restriction fragments.

To assess the impact of extending anchors to the edges of the restriction fragments, we compared loops identified using standard MACS2 ChIP-seq peaks with those based on padded anchors (as implemented in `hichipper`). (Again, for clarity, ChIP-seq peaks were used as *a priori*-defined loop anchors for the comparisons shown in Supplemental Figures 12–14 as this method of defining anchors best captures the added benefit of padding to the ends). Supplemental Figure 12 shows the number of loops identified (left) as well as the number of PETs supporting those loops (right). While we only observed a very marginal increase in the number of loops called, we observed a 35% increase in the number of PETs supporting those loops. The increased PET count is partially due to high read densities near the edges of long restriction fragments. Of note, the number of defined anchors decreases slightly (about 4% due to the merging of nearby peaks) when extending anchor loci to the ends of the restriction fragment, and the median anchor size increases by 25% (or 746 base pairs). Thus, we expect that the increased PET count will improve power in loop calling and in analyses of differential looping and other topological variation.

**Supplemental Figure 12:**
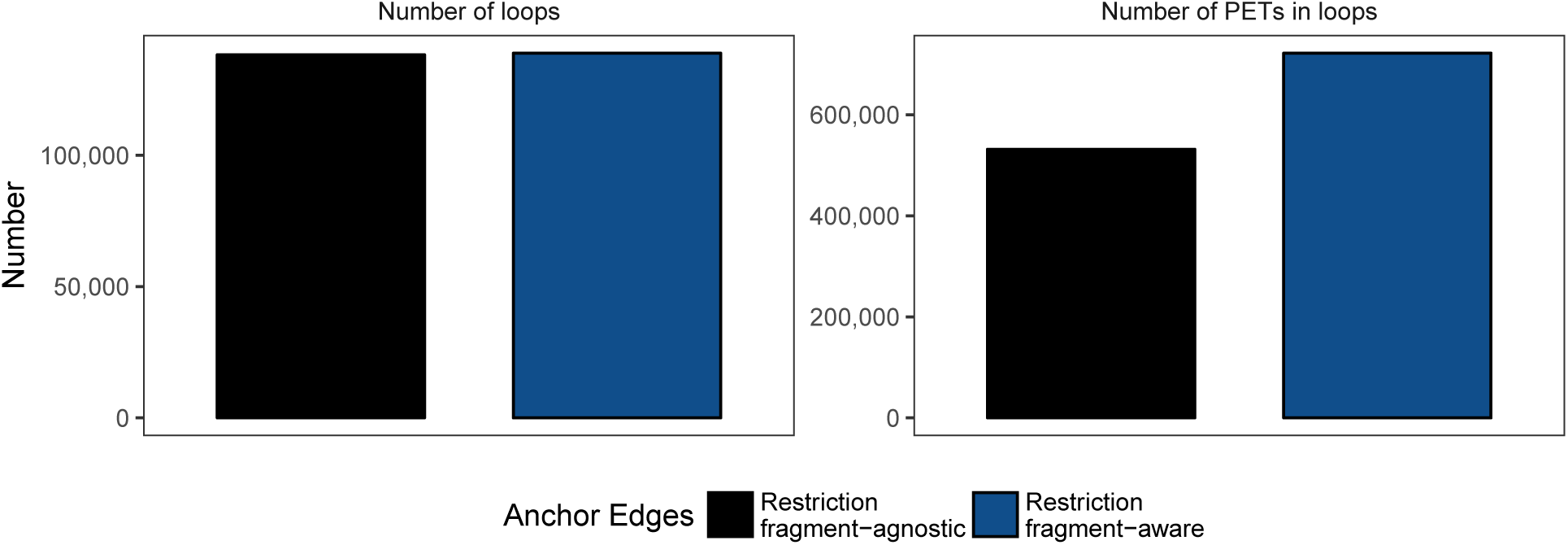
Number of loops (left) and PETs supporting loops (right) for HiChIP processing with restriction fragment-naive anchors (black), and restriction fragment-aware anchors (blue) as implemented in hichipper. For this figure, loop anchors were pre-specified with mESC SMC3 ChIP-Seq peaks to identify loops

### Loop calling

The Mango^5^ ChIA-PET pipeline implements a model for determining whether interactions are significantly stronger than the random background interaction frequency. We assessed whether this model was also suitable for HiChIP data and found it consistent with the two main assumptions of the Mango model (Supplemental Figure 13). For each assay, we defined a loop as an interaction between two loci supported by two or more PETs with a Mango *q*-value < 0.01. Using the Mango loop significance model on HiChIP data results in a large increase in sensitivity compared to ChIA-PET, due to the improved efficiency of HiChIP. A comparison of loop calls from SMC1 ChIA-PET and HiChIP in ESC cells, for example, shows a 67% increase of called loops and a 3.5-fold increase in PETs supporting those loops from 30% less sequencing (Supplemental Figure 14).

**Supplemental Figure 13:**
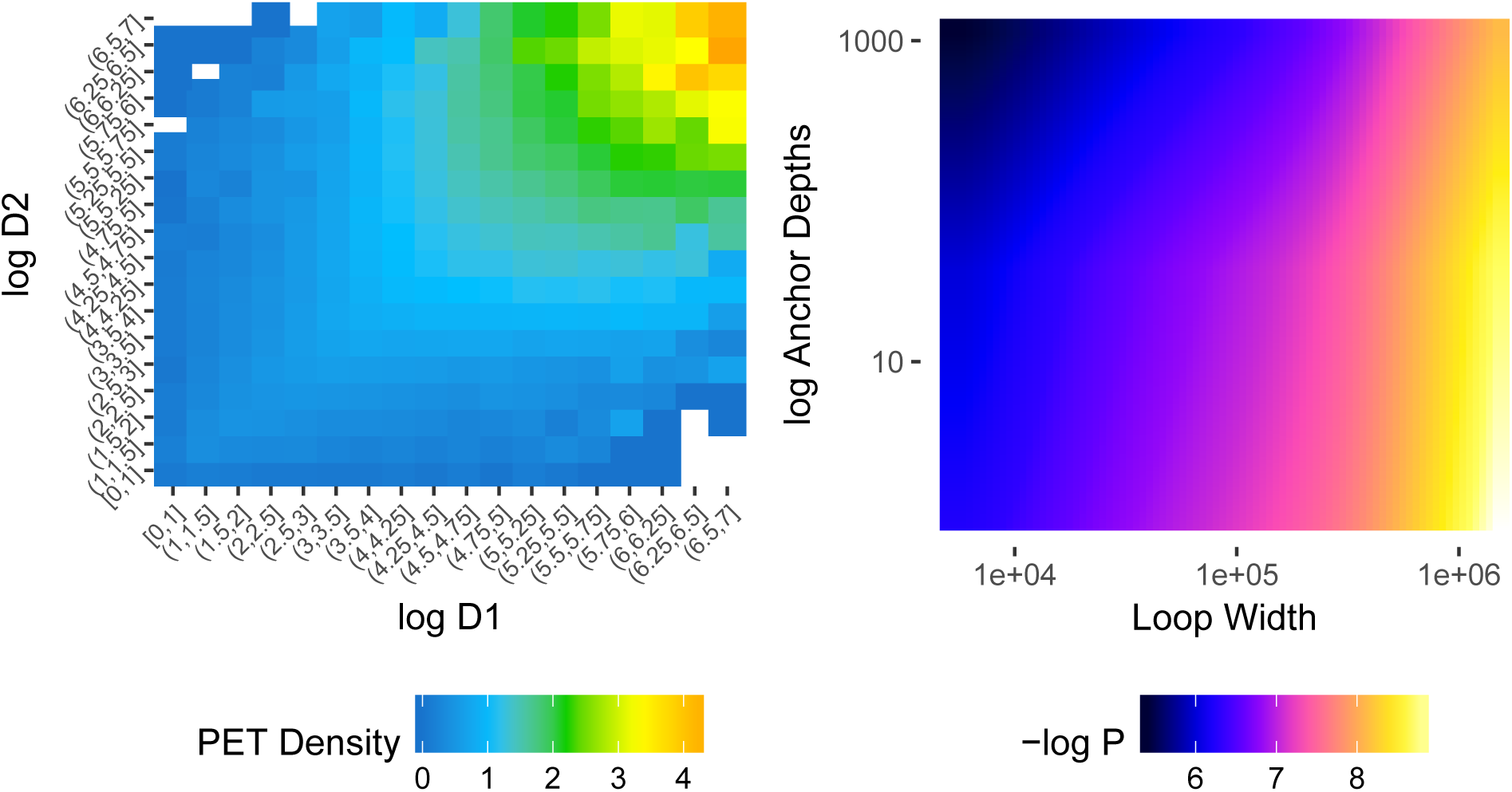
Evaluation of parametric assumptions from the Mango loop-calling model in HiChIP. (Left) PET density between unique pairs of anchors. This surface verifies the PET density (color) is roughly constant as a function of the product of the anchor depths (D1 and D2). (Right) Estimated binomial probabilities (color) as a function of genomic distance and joint anchor depth. The surface shape is similar that from ChIA-PET as depicted in Figure S1 of the Mango paper (Phanstiel *et al.*)

**Supplemental Figure 14:**
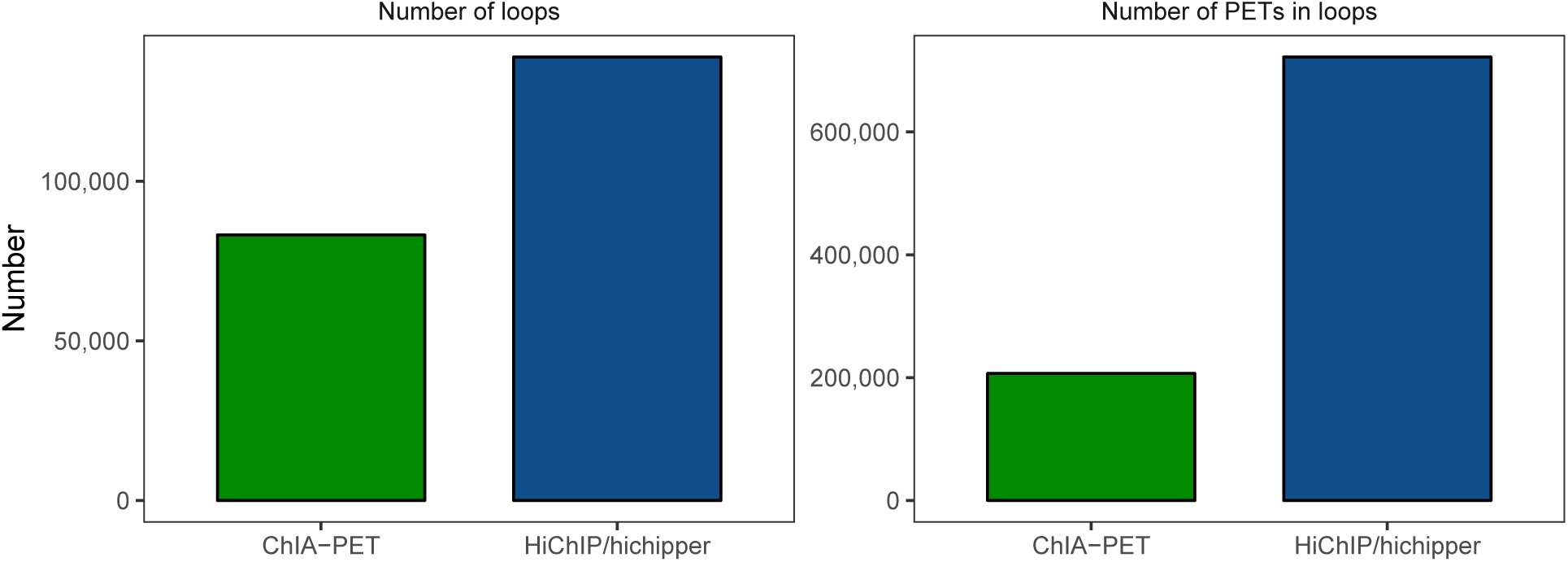
Number of loops (left) and PETs supporting loops (right) for SMC1 ChIA-PET and HiChIP of mES cells. HiChIP/`hichipper` identifies more loops and has more useful PETs than ChIA-PET, despite the lower sequencing depth. (Total HiChIP reads = 158,615,491; Total ChIA-PET reads = 221,653,525)

### Comparison with non HiChIP-specific loop-calling software

Though a variety of algorithms that identify loops from chromatin conformation capture data exist, none have been specifically designed to identify loops from assays like HiChIP with restriction enzyme digestion and chromatin immunoprecipitation. Direct use of tools such as mango^5^ and ChIA-PET2^8^, both originally designed for calling loops from ChIA-PET data and which identify peaks using a standard application of MACS2^8^, result in a high number of false positive loop anchor calls near restriction sites (Supplemental Figures 3, 8. The CHiCAGO^9^ pipeline is specifically designed for promoter capture Hi-C and restricts analysis to a pre-specified set of bait regions. We therefore compared loops identified by hichipper to those from two tools designed for HiC data - Fit-HiC^10^ and HiCCUPS (part of the Juicer^11^ suite of tools) - for two mESC SMC1 HiChIP replicates (SRR3467179, SRR3467180).

A notable difference between hichipper, Fit-HiC, and HiCCUPS is that the latter two algorithms require a loop anchor resolution specified *a priori*, consistent with standard Hi-C analysis methods that bin the genome into fixed width intervals (*e.g.* 5kb, 25kb). As the median loop anchor width from our hichipper-inferred loops was ∼2,500bp (SRR3467179: 2482bp; SRR3467180: 2539bp), we attempted to call loops using these algorithms at a pre-defined 2,500 bp resolution. However, as this resolution was not feasible with Fit-HiC ^a^ or Juicer/HiCCUPs ^b^ due to the large number of interaction pairs to be evaluated, we instead ran these tools at a 5kb resolution.

Supplemental Figure 15 shows the total loop count after running each of the three tools ^c^ on (A) the SRR3467179 and SRR3467180 replicates and (B) the loop length distribution for the SRR3467179 replicate at a 1% FDR.

**Supplemental Figure 15:**
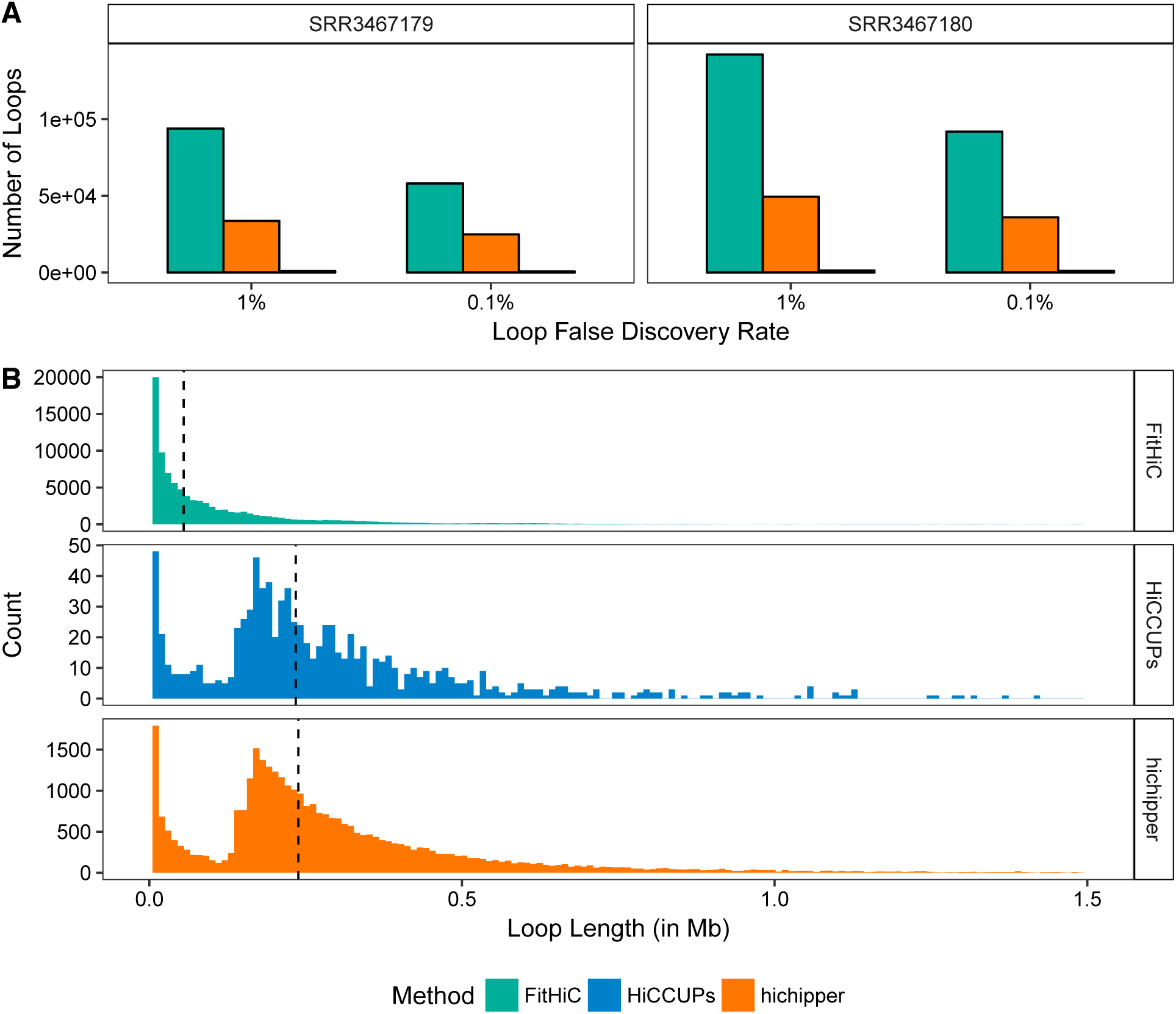
Summary of HiChIP loop calling by three different methods. (A) Number of loops called at two false discovery rates (FDR) for two mESC replicates. Fit-HiC and hichipper call greater than an order of magnitude more loops than HiCCUPs. (B) Histogram of loops called in the SRR3467179 sample at an FDR of 1%. While Fit-HiC calls approximately two-fold more loops than hichipper, nearly all of these are shorter-range interactions at less than 100kb. The distribution of loop length is very similar between HiCCUPs and hichipper. The median loop leangth for each method is indicated by the black dotted line.

As shown in Supplemental Figure 15 (A), HiCCUPs called greater than an order of magnitude fewer loops than Fit-HiC or `hichipper`. Conversely, the number of loops identified by Fit-HiC was approximately twice that of hichipper. To further examine properties of these loops, we examined the distribution of loop lengths shown in Supplemental Figure 15 (B). While the distribution of loop lengths called by hichipper and HiCCUPs is essentially identical with many loops representing interactions between loci greater than 100kb apart (shown more clearly in Supplemental Figure 16), Fit-HiC’s loop frequency decays exponentially with loop length. Moreover, while the median loop length (indicated by the black vertical line) for hichipper and HiCCUPs > 230kb, the median loop length of Fit-HiC was 55 kb. Thus while Fit-HiC calls many more loops on this HiChIP dataset, the majority of these are short-range interactions. We have found that longer-range interactions are particularly useful for many applications, including linking distal regulatory elements to target genes.

**Supplemental Figure 16:**
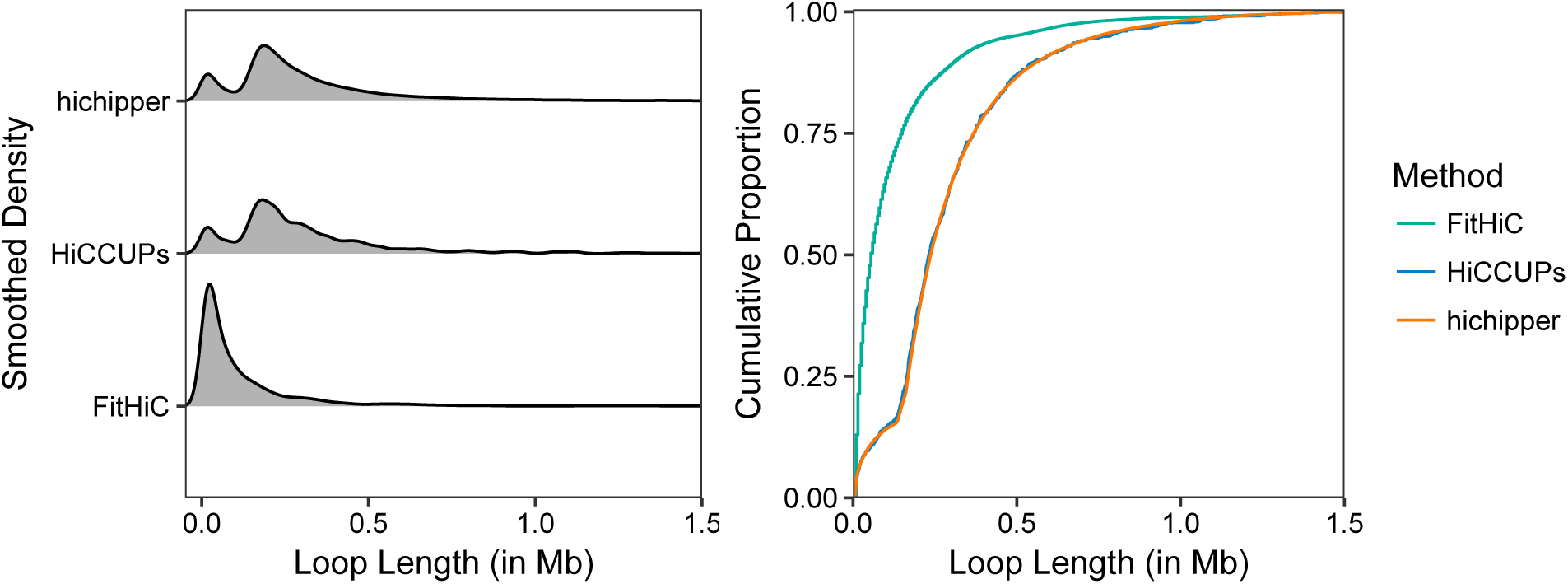
Further comparison of loop lengths called by Fit-HiC, HiCCUPs, and hichipper for the SRR3467179 sample at an FDR of 1%. Both the smoothed density (left) and cumulative distribution function (right) of loop lengths indicate that hichipper is more sensitive to detecting long-range interactions greater than 100kb.

As recent studies^12,13^ of chromatin loops have identified a preference in CTCF motif orientation, we quantified the proportion of tandem (>> and <<) and convergent (><) loops called by each tool using loops that contained a single CTCF motif at either loop anchor. The relative proportions of these loop configurations are shown in Supplemental Figure 17 where the dotted lines indicate previously reported estimates from CTCF ChIA-PET by Tang, Luo, Li et al. ^13^ of convergent and tandem loop rates (64.5% and 31.1% respectively).

**Supplemental Figure 17:**
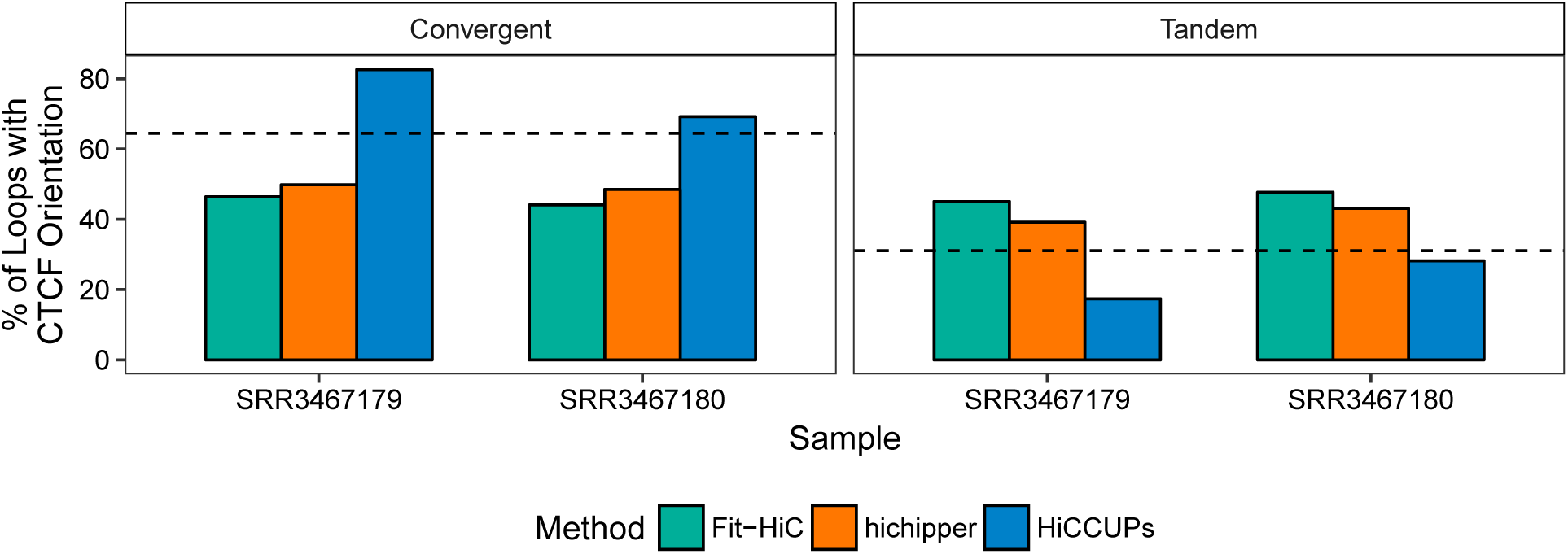
Percentage of loops with convergent (><; left) and tandem (>> or <<; right) CTCF motif orientation for loops at a FDR of 1 %.

Overall, our comparisons suggest that direct application of existing HiC tools to HiChIP data lead to either lower overall sensitivity (HiCCUPs) or lower power to detect long-range (> 100kb) interactions (Fit-HiC). `hichipper` appears to be more sensitive, while preserving the enrichment for convergent CTCF motif orientation and ChIP-seq peaks at loop anchors (Supplemental Figure 9). Inspection of interaction count matrix heatmaps further supports the validity of loops identified by hichipper with Supplemental Figure 18 showing an example loop missed by HiCCUPs, and Supplemental Figure 19 showing a loop missed by both Fit-HiC and HiCCUPs.

**Supplemental Figure 18:**
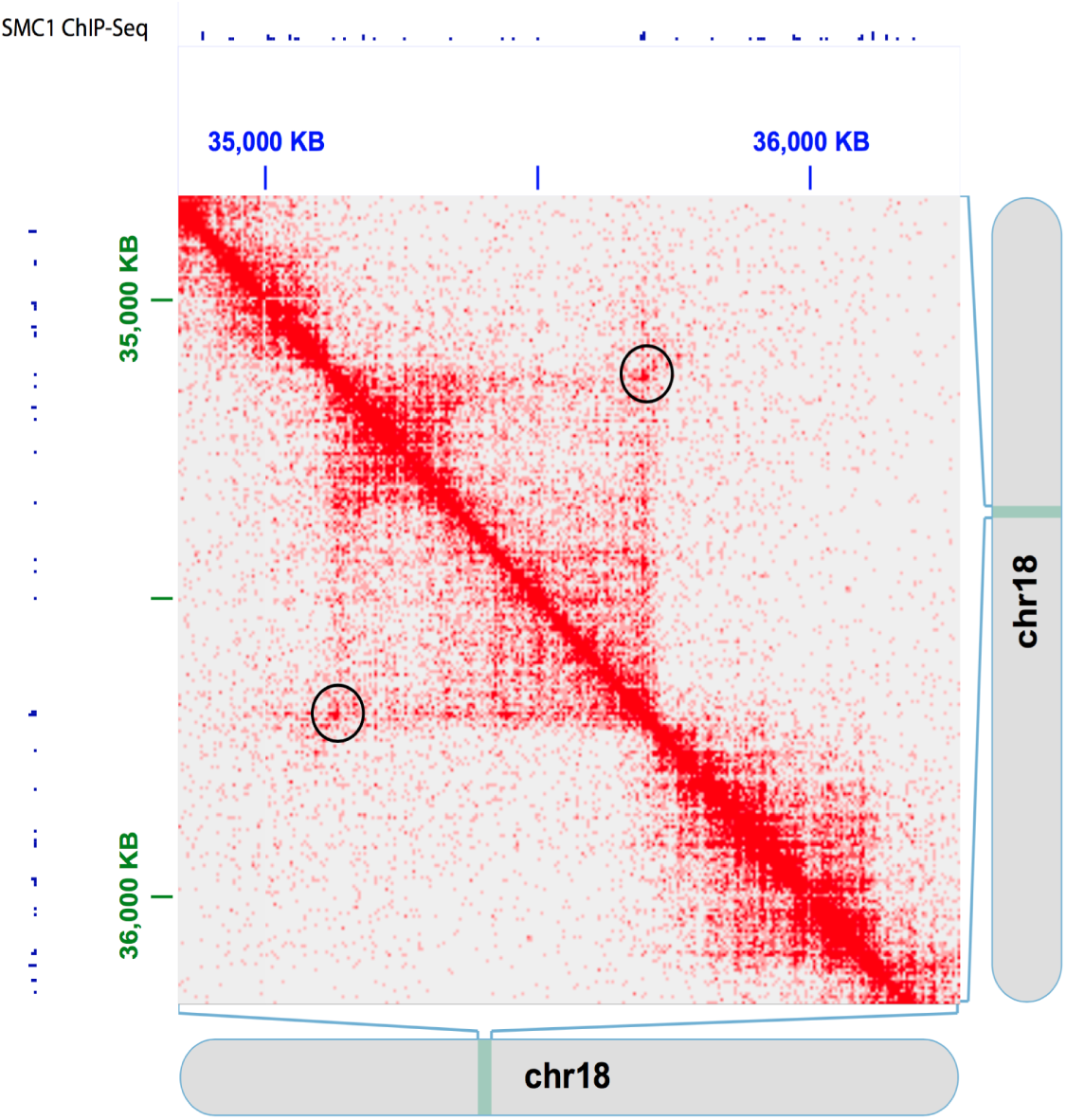
Read density map of chromsome 18 locus at 5kb resolution. The circled pixels noted in black show a long range interaction bound by two ChIP-seq peaks identified by both FitHiC and hichipper, and represents one of the few > 100 kb HiChIP interactions identified by FitHiC. No loops in the image were called by HiCCUPs.

**Supplemental Figure 19:**
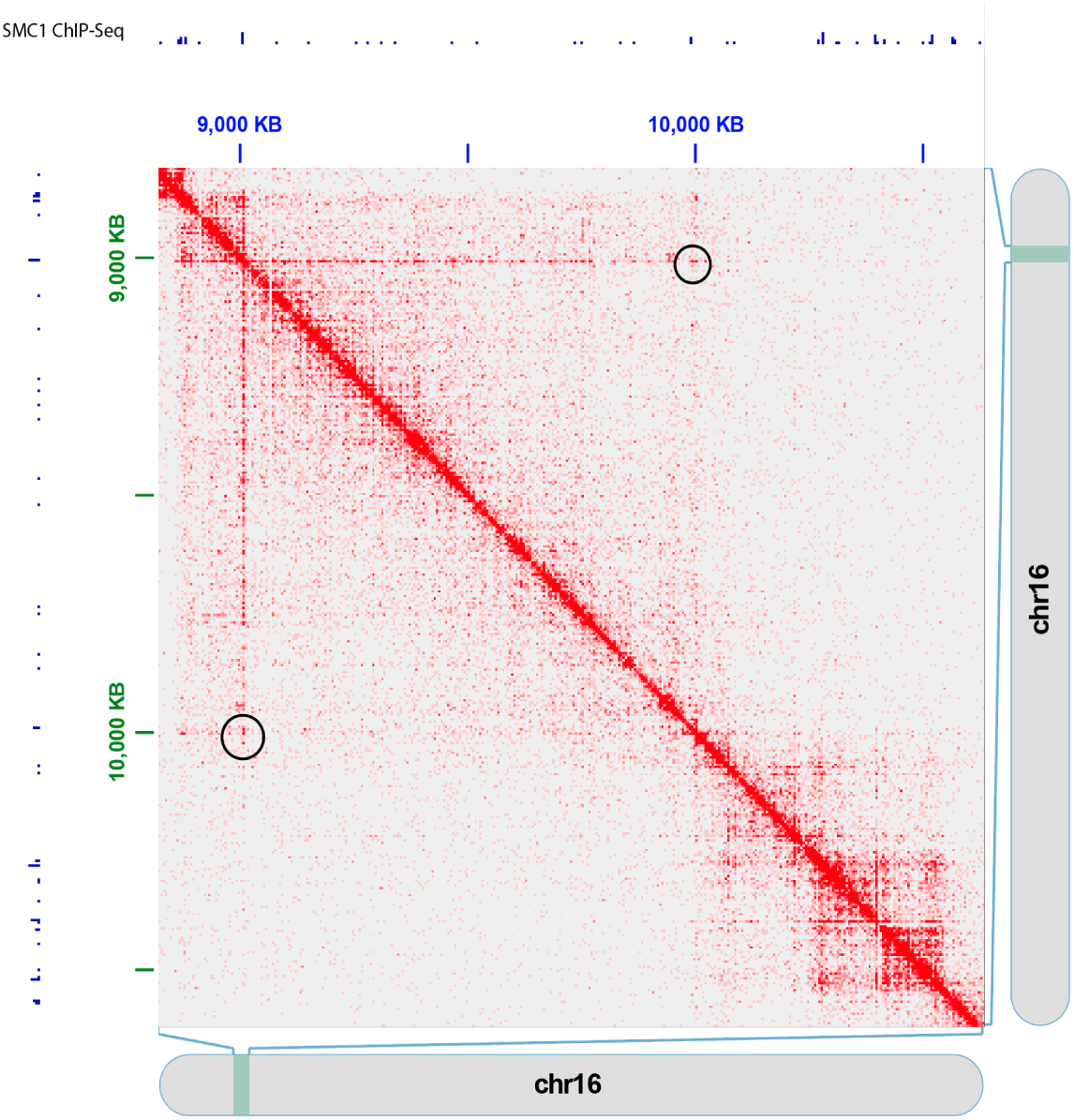
Read density map of chromosome 16 locus at 5kb resolution. The circled pixels in black represent an interaction spanning approximately 1 Mb supported by SMC1 ChIP-seq data that was called as a loop only in `hichipper`.

### Assigning putative regulatory elements to target genes

We examined the impact of loop length on downstream analyses that integrate chromatin conformation capture and genetic/epigenetic data. Specifically, a common analysis involves the assignment of epigenetic mark loci (*e.g.* H3K27ac peaks) or genetic variants (*e.g.* single nucleotide polymorphisms (SNPs) identified by GWAS) to target genes. A common approach involves assignment to the closest gene by linear distance.

**Supplemental Figure 20:**
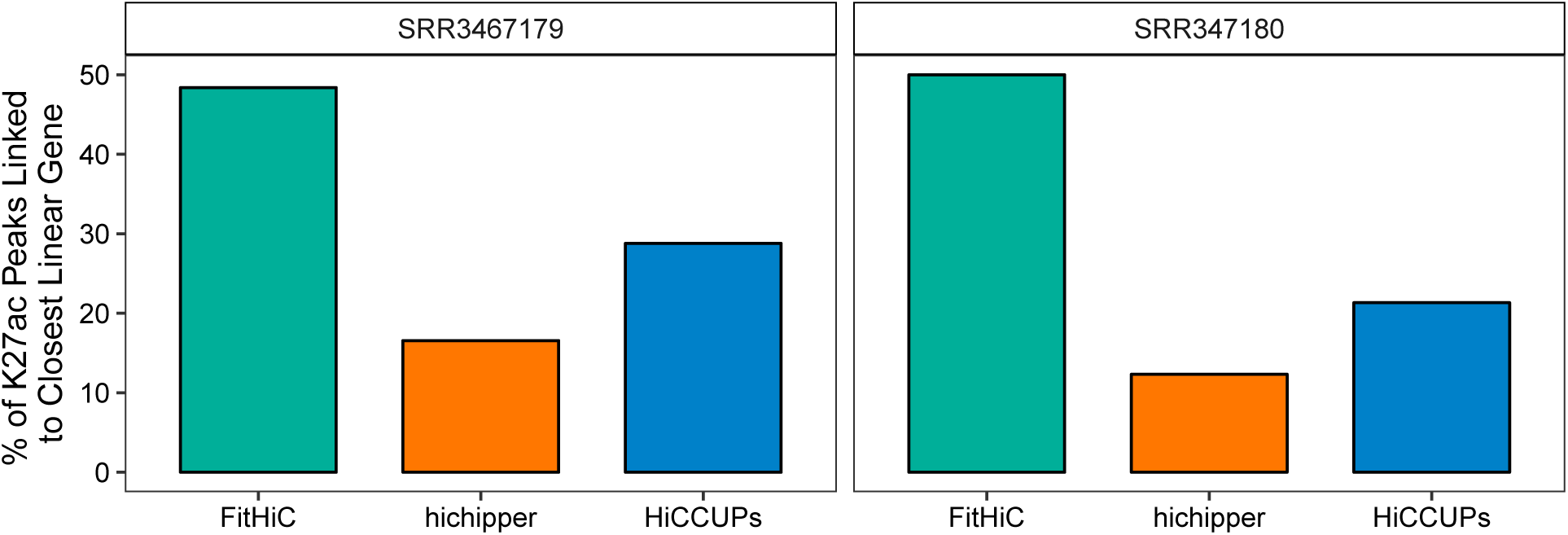
Assignment of enhancers/H3K27ac peaks to target genes. Bar heights represent the fraction of observed enhancer-promoter loops from the mESC HiChIP dataset where an enhancer is linked to the closest (linear distance) gene. The length distribution of hichipper loops is such that many enhancers are linked to genes that are not the nearest neighbor.

We called enhancer-promoter loops at a 1% FDR for each method and replicate. Less than 20% of the K27Ac marked enhancers in this set are linked to the closest TSS (Supplemental Figure 20) by `hichipper`, highlighting the value of loop data in correctly assigning target genes to regulatory elements.

### Evaluation of anchor resolution

As `hichipper` was able to infer more precise loop anchors than other methods, we examined how the increased resolution impacts the ability to identify a specific transcription start site or epigenetic mark associated with individual anchors. We considered RefSeq transcription start sites and five epigenetic features where peaks were identified from mESC ChIP-seq and DNase-seq data (Supplemental Figure 21 - left). We examined the ambiguity associated with identifying a specific feature as a function of anchor resolution (right). To compute this metric, we padded each peak or transcription start site to represent a loop anchor centered on this feature. We then counted the average number of overlaps with other features in the set. For example, at a resolution of 0 base pairs, each feature only overlaps itself, leading to an ambiguity of 1. As the padding (anchor size) increases, we are increasingly likely to overlap with other features making the assignment ambiguous (>1).

**Supplemental Figure 21:**
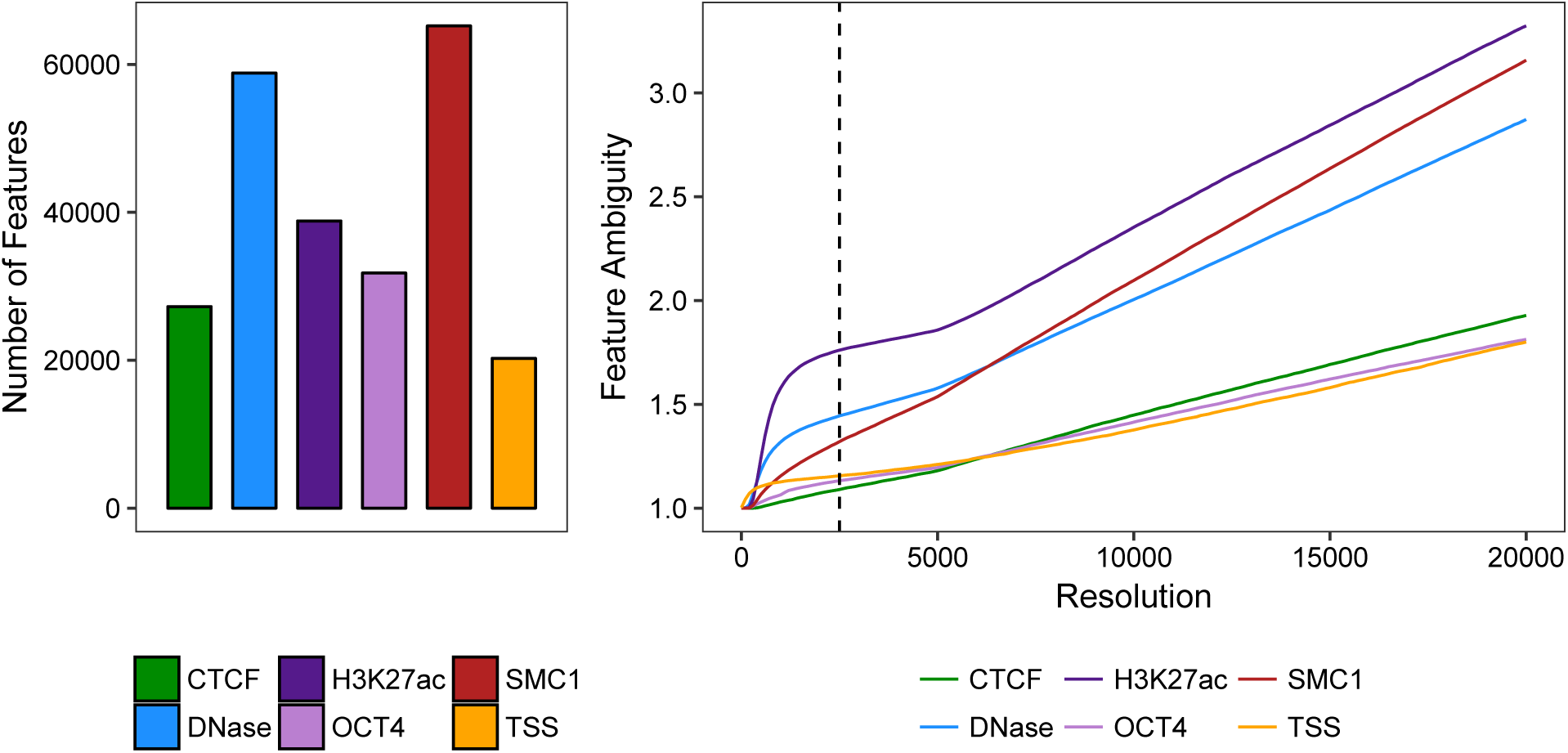
(Left) The number of peaks and transcription start sites used for comparison. (Right) To assess the impact of high-resolution loop anchor calling, the mean feature ambiguity is plotted against the loop anchor resolution. The dotted vertical line indicates the median resolution of loop anchors from hichipper.

At a resolution of 20kb, a common Hi-C bin width, a mean ambiguity for DNase peaks is 2.5. As such, loop calls at 20kb resolution seldom resolve individual open chromatin loci. However, a 2.5kb resolution, the median resolution for loop anchors called by `hichipper`, enables the assignment of many single genetic and epigenetic features to individual loop anchors. We expect that the resolution of `hichipper` loop anchors will prove useful, as many chromatin capture analyses are designed to resolve enhancer-promoter contacts or assign putative functional targets for specific transcription factors.

### Library quality metrics

`hichipper` generates quality control metrics that can be used to assess HiChIP libraries, including the proportion of long range interactions (related to proximity ligation efficacy) and the proportion of reads mapping in `hichipper`-defined anchors (related to ChIP efficacy). We provide example quality-control reports for a variety of successful and unsuccessful HiChIP experiments at the hichipper website (http://aryeelab.org/hichipper). These will serve as a reference for new adopters of the HiChIP assay, enabling quality comparisons of user-generated libraries to `hichipper` QC reports for existing data.

### Availability

Source code and documentation for `hichipper` is available at http://aryeelab.org/hichipper. The software is also distributed as a PyPi package, available at https://pypi.python.org/pypi/hichipper/.

## Footnotes

a The resolution limitation relates to the number of interaction pairs exceeding an `R/data.table` limit of 2^31^. The specific error message returned in the **R** console was the following: Join results in more than 2^31^ rows (internal vecseq reached physical limit). After further examination of the Fit-HiC source code, the software attempts to create a `possiblePairsPerDistance` object using the `data.table` package, which causes the error at the 2.5kb resolution. For example, the length of chromosome 1 in the mm9 mouse genome build is 197 Mb, resulting in ((*n*\**n* (*n* − 1))*/*2 > 2^31^*possiblepairs*; *n* = 197, 000, 000*/*2, 500 (all combinations of the chromosome 1 bins at 2.5kb resolution).

b The specific error message returned from the **java** console was the following: `Discarding invalid configuration: Configres: 2500 peak: 4 window: 1 fdr: 10% radius:` 1 This same error message occurred after attempting to run HiC-CUPs with both the default (unspecified) and suggested (manually specified `-f 0.1 -w 7 -p 4`) parameters. Though we confirmed that the 2,500bp zoom option present in the .hic file, previous uses of the HiCCUPs algorithm have called loops at a 5kb and 10kb resolution^12^

c HiCCUPs was run with the `--ignore-sparsity` flag. Fit-HiC was run with default parameters.

## References

(1) Mumbach, M. et al. Nature Methods 13, 919–922 (2016).

(2) Servant, N. et al. Genome Biol. 16, 259 (2015).

## References

Kagey, M.H. et al. Mediator and cohesin connect gene expression and chromatin architecture. Nature 467, 430–435 (2010).

Dowen, J.M. et al. Control of cell identity genes occurs in insulated neighborhoods in mammalian chromosomes. Cell 159, 374–387 (2014).

Mumbach, M.R. et al. HiChIP: efficient and sensitive analysis of protein-directed genome architecture. Nat Methods 13, 919–922 (2016).

Langmead, B. & Salzberg, S.L. Fast gapped-read alignment with Bowtie 2. Nat Methods 9, 357–359 (2012).

Phanstiel, D.H., Boyle, A.P., Heidari, N. & Snyder, M.P. Mango: a bias-correcting ChIA-PET analysis pipeline. Bioinformatics 31, 3092–3098 (2015).

Servant, N. et al. HiC-Pro: an optimized and flexible pipeline for Hi-C data processing. Genome Biol 16, 259 (2015).

Zhang, Y. et al. Model-based analysis of ChIP-seq (MACS). Genome Biol 9, R137 (2008).

Li, G., Chen, Y., Snyder, M.P. & Zhang, M.Q. ChIA-PET2: a versatile and flexible pipeline for ChIA-PET data analysis. Nucleic Acids Res 45, e4 (2017).

Cairns, J., et al. CHiCAGO: robust detection of DNA looping interactions in Capture Hi-C data.Genome Biol 17, 127 (2016)

Ay, F., Bailey, T., & Noble, W. Statistical confidence estimation for Hi-C data reveals regulatory chromatin contacts. Genome research 24, 999–1011 (2014).

Durand, N. et al. Juicer provides a one-click system for analyzing loop-resolution Hi-C experiments.Cell systems 1, 95–98 (2016).

Rao, S. et al. A 3D Map of the Human Genome at Kilobase Resolution Reveals Principles of Chromatin Looping. Cell 7, 1665–1680 (2014).

Tang, Z. et al. CTCF-mediated human 3D genome architecture reveals chromatin topology for transcription. Cell 7, 1611–1627 (2015).

